# Drawing the Borders of Olfactory Space

**DOI:** 10.1101/2020.12.04.412254

**Authors:** Emily J. Mayhew, Charles J. Arayata, Richard C. Gerkin, Brian K. Lee, Jonathan M. Magill, Lindsey L. Snyder, Kelsie A. Little, Chung Wen Yu, Joel D. Mainland

## Abstract

In studies of vision and audition, stimuli can be chosen to span the visible or audible spectrum; in olfaction, the axes and boundaries defining the analogous odorous space are unknown. As a result, the population of olfactory space is likewise unknown, and anecdotal estimates of 10,000 odorants have endured. The journey a molecule must take to reach olfactory receptors (OR) and produce an odor percept suggests some chemical criteria for odorants: a molecule must be 1) volatile enough to enter the air phase, 2) non-volatile and hydrophilic enough to sorb into the mucous layer coating the olfactory epithelium, 3) hydrophobic enough to enter an OR binding pocket, and 4) activate at least one OR. Here, we developed a simple and interpretable quantitative model that reliably predicts if a molecule is odorous or odorless based solely on the first three criteria. Applying our model to a database of all possible small organic molecules, we estimate that over 30 billion possible compounds are odorous, 6 orders of magnitude larger than current estimates of 10,000. With this model in hand, we can define the boundaries of olfactory space in terms of molecular volatility and hydrophobicity, enabling representative sampling of olfactory stimulus space.

---

The number of molecules humans can smell is disputed, with published estimates ranging from 10,000^1^ to infinitely many^2^. Chemical space is vast, and we cannot resolve this dispute until we define the subset of chemical space that has an odor.Without defining the boundaries of this space, we cannot know whether previous research has adequately sampled from it, nor understand how the brain represents it, nor conduct a rational search for novel odorants within it.

To act as an odorant, a molecule must complete a transport process to reach olfactory receptors (OR) and activate one or more ORs (Fig. 1a). Viewing the journey of an odorous molecule to an olfactory receptor (OR) as a mass transport problem^3^ points to the types of molecular criteria we must consider (e.g. vapor pressure, hydrophobicity); but because no previous studies have both proposed and empirically tested a biological theory of the chemical criteria for an odorant, the relative importance, precise limits, and interactions of these constraints remain undefined.

**Figure 1:**
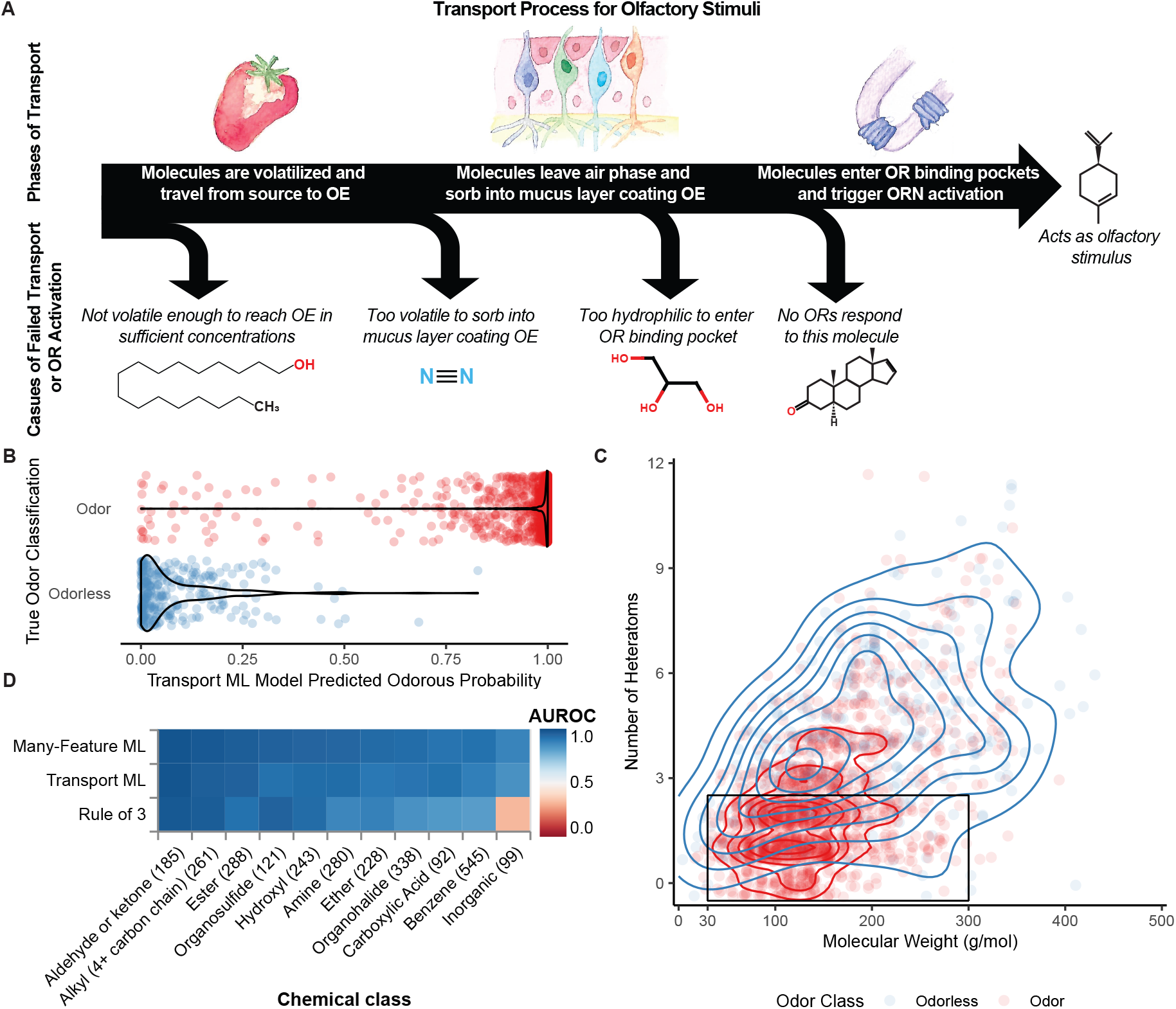
A machine learning model can accurately classify molecules as odorous or odorless based only on transport features. (**A**) Schematic of the transport process molecules must complete to act as olfactory stimuli. To elicit an odor, molecules must reach the olfactory epithelium (OE), enter olfactory receptor (OR) binding pockets, and trigger OR neuron (ORN) activation. (**B**) Transport-feature ML model-generated odorous probabilities for all molecules in the dataset. Each dot represents one molecule colored by the ground truth, and the width of the violin plot is the density of molecules at a given prediction value. (**C**) Density of odorous and odorless molecules in transport space defined by molecular weight and number of heteroatoms. Each successive contour line indicates a step increase in density (red = 0.05%, blue = 0.01%). Molecules have a discrete number of heteroatoms, but are jittered along the y-axis to better show density. Plotted within the black box, molecules that obey the “rule of three” with fewer than 3 heteroatoms and molecular weight between 30 and 300 g/mol are generally odorous. Relative performance of models depends on functional group/chemical class. (**D**) Heatmap of mean AUROC generated by four models for molecules of common chemical classes (number of matching molecules in parentheses).

## Building an odor classification model

We set out to learn which chemical criteria separate odorous from odorless molecules by building the simplest accurate model from a diverse pool of molecules and chemical features. A model using only three features that dictate transport capability (boiling point, vapor pressure, and octanol/water partition coefficient “log P”) reliably classifies molecules as odorous or odorless (Fig. 1b). If odor classification can be explained by so few molecular properties, why were the classification rules not previously known? The likely reason is that available data are both noisy and poorly curated; we needed to gather odor classification data from multiple sources and correct errors in the data – both in transport features and odorous/odorless labels – before the three-parameter transport model matched the performance of more complicated models.

The theory that transport properties define odor space was previously proposed almost 40 years ago by Boelens^4^, but the manuscript provided no empirical evidence for the claim. More recent efforts to predict odorous/odorless classification of molecules have relied on existing flavor and fragrance databases^5–7^; the biased scope of molecules and classification errors in these databases constrain the success of these models. A recent neural network-trained odorous/odorless classification model advertised high accuracy, but we found that its classification accuracy was poor when tested on our cleaned dataset (AUROC = 0.626; Extended Data Fig. 1)^6^.

Ultimately, we generated a large and chemically diverse dataset of over 1,900 molecules, classified as odorous (84%) or odorless (16%) through a combination of literature-and web-scraping, human discrimination tasks, and chemical analysis. We set aside in advance a test set of 30 molecules with high-confidence classifications (classified by human subjects and confirmed through chemical analysis) to measure final model performance. The remaining molecules formed our training set, and we applied machine learning (ML) algorithms to train odor classification models. Our models represent molecular structure as a vector of physicochemical features (Dragon v6, Talete; EPI Suite, U.S. EPA), and calculate a probability that the molecule is odorous; all code used to generate models and figures is publicly available (https://github.com/emayhew/OlfactorySpace).

We optimized ML models and measured performance using the area under the receiver operating characteristic curve (AUROC), a metric for which 1.0 represents perfect classification and 0.5 represents chance-level classification accuracy. We achieved near-perfect AUROC values in cross-validation with several algorithms when paired with a synthetic minority oversampling technique (SMOTE) to address the imbalance in odorous:odorless training examples. When applied to the test set, a simple three-parameter ML model separates odorous from odorless molecules with no mistakes (AUROC = 1.0). Our test set was designed to maximize chemical diversity within safety and availability constraints, so strong predictive performance on this held-out set suggests that the model will generalize well to new molecules.

We next compared the performance of our transport ML model to a “many-feature” ML model trained with over 3700 physicochemical features. Even when regularization is applied to reward use of few, highly relevant features to generate predictions, adding features does not improve classification accuracy (AUROC = 0.94). This finding supports the theory that transport-capability is what determines whether a molecule is odorous or odorless.

Our transport ML model achieves high accuracy with few features, but experimental values of boiling point, vapor pressure, and log P are unavailable for many molecules. We also developed a simple rule of thumb that sacrifices some accuracy but can be applied knowing only the molecular formula (e.g. C10H12O2): the “rule of three” states that molecules with molecular weight between 30 and 300 Da and with fewer than 3 heteroatoms are generally odorous (Fig. 1c). In our dataset, 96% of the molecules that meet these criteria are odorous (positive predictive value); these criteria capture 68% of odorous molecules (sensitivity) and exclude 84% of odorless molecules (specificity).

Next, we asked if the relative accuracy of the transport ML, many-feature ML, and “rule of three” models varied by chemical class. Fig. 1d shows the test set AUROC for common chemical classes, averaged over 80 models trained on randomized train/test splits. All models achieved strong predictive accuracy for alkanes, alcohols, and carbonyl-containing molecules, but only ML models accurately classified organohalides. The “rule of three” underperforms on inorganic compounds because while molar mass is a good proxy for volatility of organic compounds (e.g. propane: MW = 44 g/mol, BP = -42 °C; butane: MW = 58 g/mol, BP = -0.5 °C), this relationship does not hold for inorganic compounds (e.g. NaCl: MW = 54 g/mol, BP = 1465°C). The strong performance of the transport ML model independent of chemical class supports the reliability of the model across common classes of molecules.

### Importance of high-quality experimental data

Building a high performing transport model required correction of two major sources of error: boiling point values and odor classifications. Boiling points (BP) reported in compendia or by chemoinformatic software are commonly estimated from chemical formula^8,9^, but these estimates are error-prone (median absolute error of 55 °C via Burnop method and 37 °C via Banks method) (Fig 2a). Using estimated BP values in an otherwise identical ML transport model worsens the classification performance (Fig. 2B-C: cross-validation AUROC = 0.93 (B), 0.95 (C); error rate = 6.0% (B), 4.5% (C)) compared to our final ML transport model (Fig. 1B: CV AUROC = 0.99, error rate = 2.6%). We replaced estimated BP with experimentally-derived values (EPI Suite, U.S. EPA) in our dataset and used estimates only when no experimental values were available. Experimental values should thus be used to generate high confidence odorous/odorless predictions, though estimated values may be adequate where a higher error rate is acceptable.

**Figure 2:**
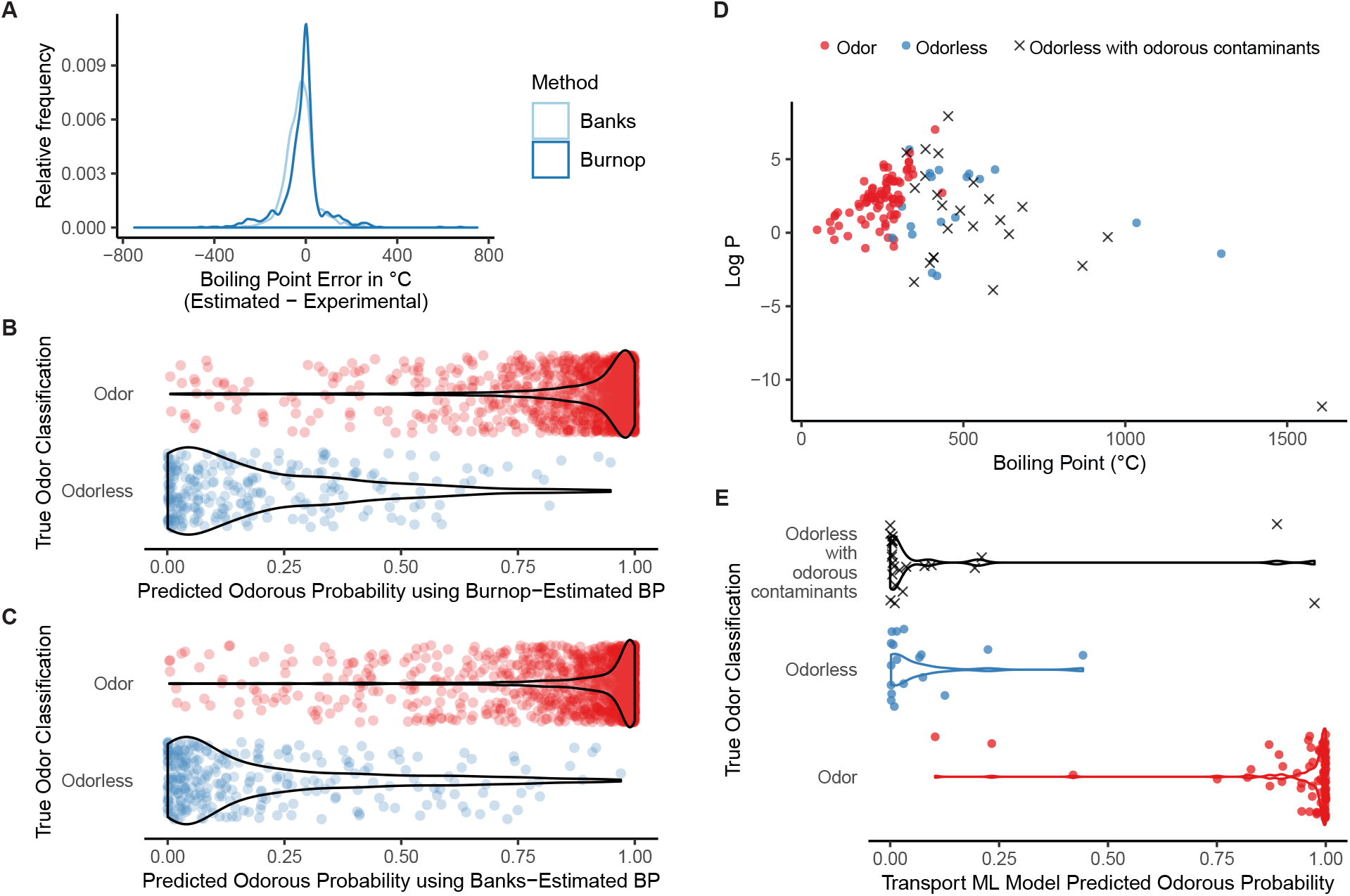
Inaccuracies in data impact model performance. (**A**) Difference between experimentally determined boiling point (BP) values and BP values calculated using the Burnop^9^ and Banks^8^. (**B** - **C**) Odor classification predictions by transport-feature ML models using only estimated BP values calculated by the (B) Burnop or (C) Banks method. (**D**) Human subject-classified molecules in transport space defined by BP and log P. Many clearly non-volatile molecules were initially classified as odors due to odorous contaminants. (**E**) Transport-feature ML model odor predictions for human subject-classified molecules. Chemical compounds that are odorless but had odorous contaminants are correctly predicted to be odorless by the model.

The second major source of error that we rectified is the odor classifications themselves. ML models can tolerate some noise in the training data, but inaccuracies in the test set can be more costly; specifically, predictive performance is bounded from above by mis-labeled data in the test set^10^. To ensure our test set was composed of accurately classified molecules, we supplemented our literature and web-scraped data with 128 molecular odor classifications (111 odorous, 17 odor-less) generated through human psychophysics experiments. We analyzed all 111 compounds with a human-detectable odor using paired GC-MS and GC-O as a quality control (QC) measure to identify cases in which impurities, rather than the target compound, were responsible for the odor detected by human subjects.

We found that 22% of molecules classified as odorous were actually odorless compounds contaminated with odorous compounds, despite high nominal purity ratings from vendors (Fig. 2D). Had we not performed this QC, we would have falsely believed that model performance was poor (pre-corrected Transport ML AUROC = 0.81). In fact, most disagreements between our model’s predictions and pre-QC classifications were due to the model correctly identifying misclassifications in the dataset (Fig. 2E).

Chemical compounds are common stimuli in olfaction research, but the impact of impurities on data is rarely discussed. Odor detection thresholds vary by many orders of magnitude across molecules, so even high purity (e.g. 99%) is insufficient grounds to consider odor to be driven entirely by the dominant molecular species^11^. Addressing the impact of contaminants on our data was thus vital for accurately measuring model performance.

### Enumerating odor space

The dispute over the size of odor space has endured in the field because the criteria for odorous molecules were not rigorously defined. We now have a tool to address this debate: a simple quantitative model based on the well-developed field of physical transport that makes highly accurate odor classification predictions and generalizes to new molecules. Chemical space is vast, but it can be enumerated. Applying graph theory and rational chemical constraints (e.g. bonds per atom, bond angles, ring strain), Ruddigkeit et al.^12^ developed GDB-17, a database of 166 billion unique molecules with heavy atom count (HAC) of 17 or fewer. While it excludes some known odorants (e.g. silica-containing molecules), the composition of this database of small organic molecules (limited to atoms C, H, N, O, S, halogens; HAC *≤* 17; unstable structures eliminated), makes it well-matched to the goal of finding new odorous molecules.

Of the 166 billion molecules in GDB-17, our transport model predicts 40 billion molecules will be odorous. This is a conservative estimate of the size of odor space because we know the odorous range extends beyond HAC 17 and that GDB-17 does not include all possibly odorous chemical structures. To approach a more realistic estimate, we 1) set an upper limit of 21 heavy atoms^5^, 2) fit a logistic regression model to predict the rate of decline in the proportion of odorous molecules with increasing HAC (Fig 3a), 3) assumed HAC 18-21 have at least as many molecules each as HAC 17, and 4) applied these extrapolated odorous proportions to HAC 18-21 (Fig 3b,c). This approach yields an estimate of 69 billion odorous molecules.

**Figure 3:**
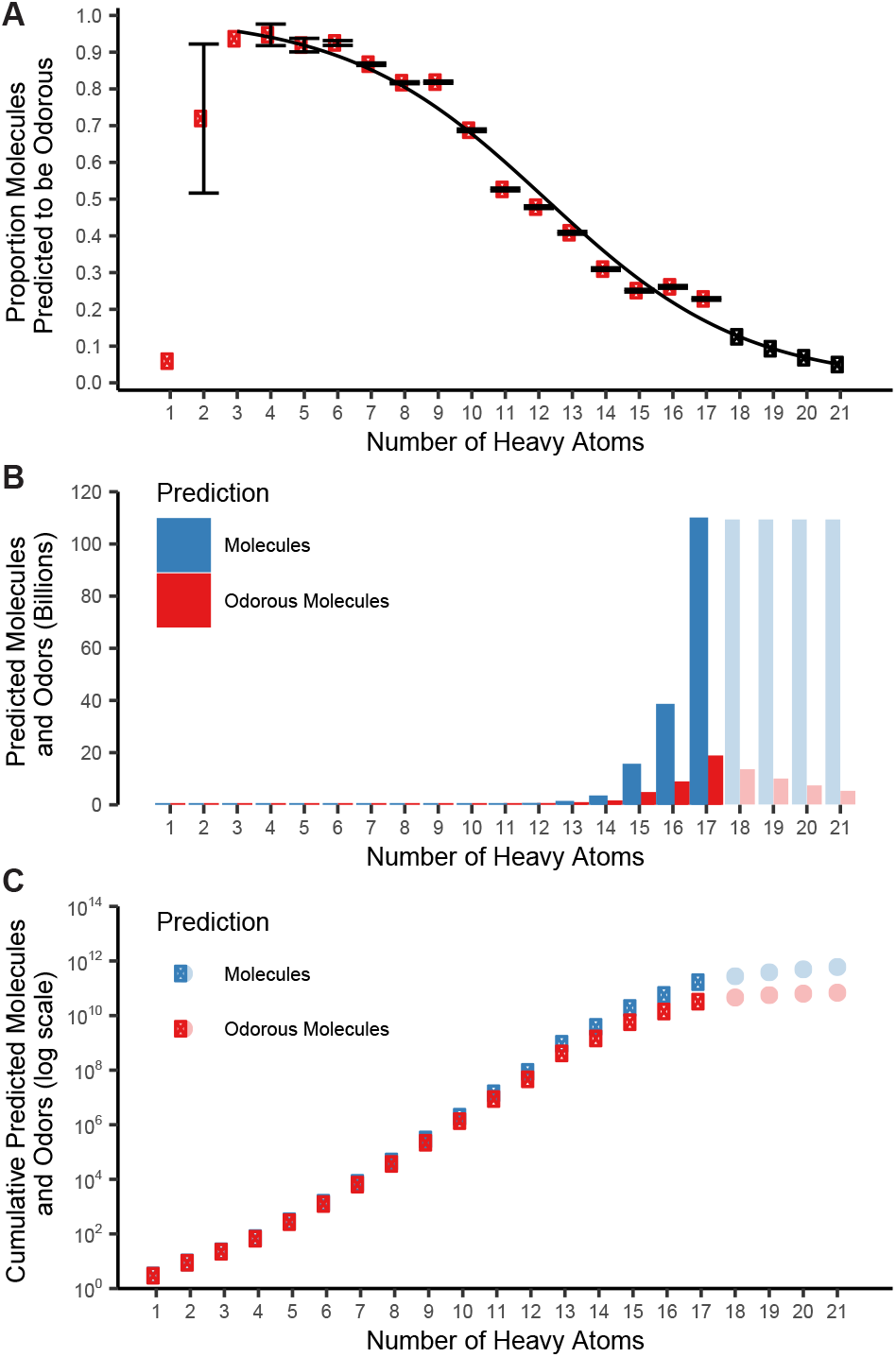
The transport model can be used to predict the population of odor space. (**A**) Proportion of molecules predicted by transport ML model to be odorous as a function of heavy atom count (HAC). Red circles show the mean probability generated for HAC tranches from the GDB database^12^ with standard error bars; hollow black circles show the projected odorous probability generated by the logistic regression fit plotted in black. (**B**) Estimated number of possible molecules and predicted odorous molecules from the GDB databases as a function of HAC. Lighter shaded bars indicate HAC values beyond GDB-17; as a conservative estimate, the number of possible molecules with HAC 18-21 was set to equal that of HAC 17. (**C**) Cumulative estimates of possible molecules and odorous molecules with increasing HAC on a logarithmic scale. The red datapoint at HAC 17 reflects our conservative estimate of 33 billion odorous molecules; the light red datapoint at HAC 21 reflects our less-conservative estimate of 62 billion odorous molecules.

The laws of physical transport upon which our model is built apply to all molecules. While most datasets used to study this question come from chemically biased flavor and fragrance databases, we intentionally purchased and classified the most chemically diverse set of molecules possible within the bounds of safety and availability. Because transport-capable molecules were almost universally odorous in our chemically diverse test set, we expect the model will successfully extrapolate to all of GDB-17. Even if we restrict our search to molecules in GDB-17 with neighbors in our training data (Tani-moto similarity ¿ 0.4), we still find 10^7^ unique predicted odorants (Extended Data Fig. 2), a value 3 orders of magnitude larger than the commonly cited estimate of 10,000 odorants^1^.

### Mapping odor space

Any database or catalog of purchasable odorous molecules is dwarfed by the scale of odor space. For example, the Sigma-Aldrich Flavor and Fragrance Catalog typically lists only 1,600 molecules. Many densely populated regions of odorous chemical space (Fig. 4A, interactive version available at https://odormap.pyrfume.org) are sparsely represented by known odorants (Fig. 4B). Our map of the odorous region of chemical space can identify new likely odorants (Fig. 4C, 4E) and be used to filter out unlikely odorants (Fig. 4D). The sheer number of as-yet-unsynthesized odorous molecules is striking and there are whole classes of odorous molecules that have not been synthesized and whose odor characteristics are to date unknown.

**Figure 4:**
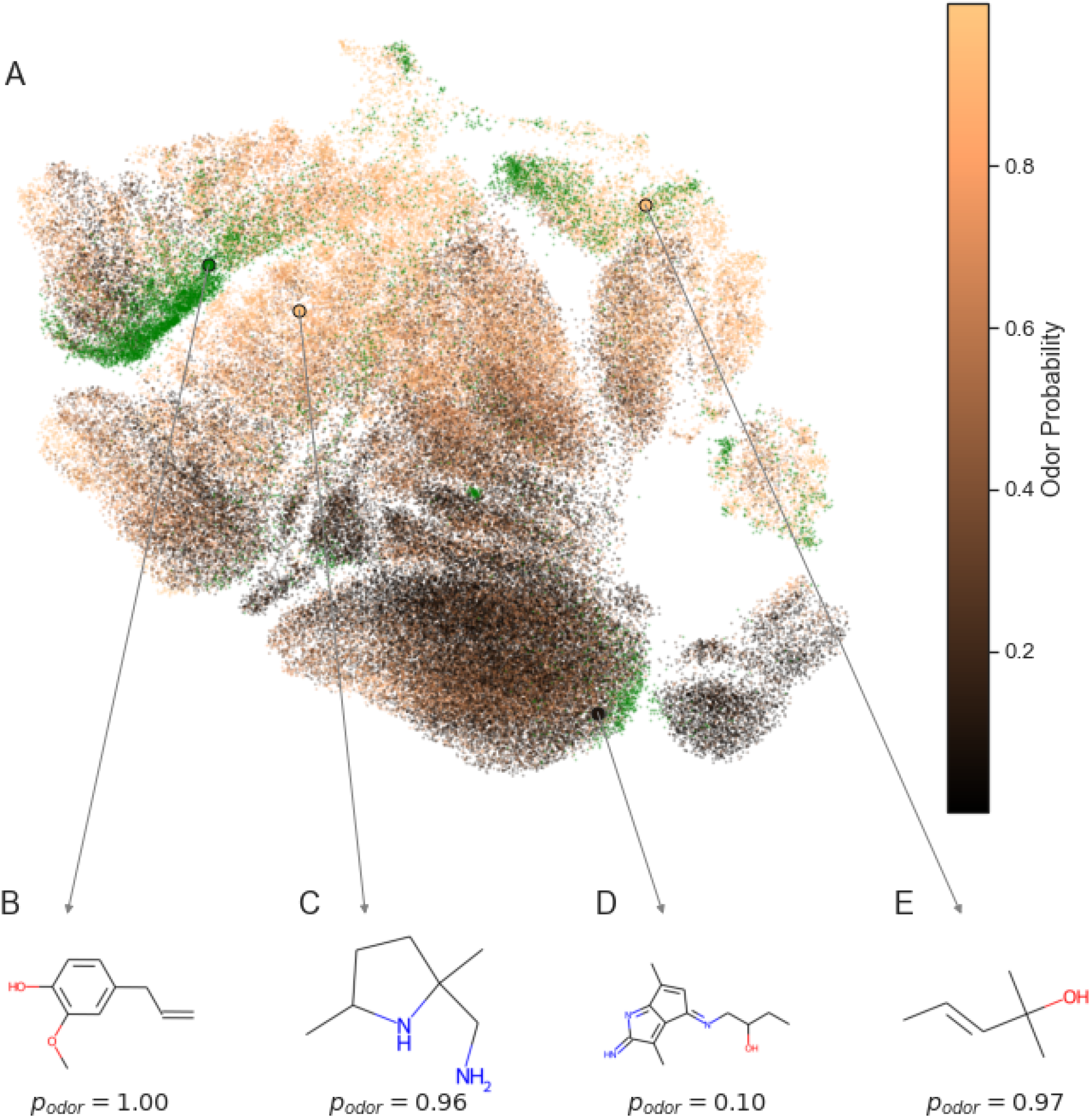
Visualization of olfactory space highlights understudied regions. (**A**) Uniform Manifold Approximation and Projection (UMAP) plot of known odorous molecules (green) and possible molecules from GDB-17 colored by their transport ML-predicted odorous probability. Many regions dense with probable odors are sparsely represented by known odors. (**B**) Eugenol, a known odorant. (**C-E**) Example molecules from GDB-17 and their transport ML-predicted probability of being odorous (*p*_*odor*_).

Drawing the borders of olfactory space is largely a matter of understanding physical transport. Nearly all molecules that can physically enter a receptor binding pocket have an odor. Although there are some clear exceptions, for some individuals, such as androstenone^13^, few molecules appear to be odorless simply because there is no sufficiently sensitive receptor. Another piece of evidence is that the odor status of a molecule can change when environmental conditions change its transport-capability: methane is odorless at standard pressure but reportedly smells of camphor when given to divers at 13 atmospheres^14^. The receptor repertoire over the course of human evolution could not have evolved to detect this molecule, yet this repertoire maintains enough general sensitivity to – once given access to it – detect it nonetheless.

We aimed to collect a chemically diverse dataset, but were limited by the types of molecules for which odor classifications are published, availability of molecules for purchase, and safety of molecules for human testing. There may be classes of odorous molecules that were not included in our training data and therefore not represented by our model. Our model was trained on human-made odor classifications, and so the resultant model should be understood to draw boundaries of human olfactory space. While we think it likely that transport features delineate olfactory space across species, differences in mucosal properties and receptor populations may shift these boundaries.

Our model answers the question of what makes a molecule odorous, but many questions remain. Our estimate of the number of possible odorous molecules cannot resolve the number of discriminable odor percepts^15–17^; many of these predicted odorants may have indistinguishable odor percepts, and odorant mixtures may produce percepts that are distinct from that of any single odorant. Everything we know about odorants is derived from a tiny subset of all volatiles—a catalog of all volatiles present in foods represents less than 0.000003% of the molecules we can smell. This model invites researchers into the unknown, providing a map to new regions of odor space and the means to representatively sample it.

## Acknowledgements

The authors acknowledge support from the George Preti Research Support Core for Analytical Chemistry and would like to thank George Preti, Katherine A. Prokop-Prigge, Bruce Kimball, Kai Zhao, and Gary K. Beauchamp for comments and discussions.

## Author contributions

C.J.A., J.M.M., L.L.S., and C.W.Y. collected human behavioral data. L.L.S., K.A.L., and E.J.M carried out quality control using GC-O. E.J.M., C.W.Y., K.A.L., R.C.G., and J.D.M. curated external data. B.K.L. validated the models and analyzed the domain applicability. C.W.Y and C.J.A conducted initial data cleaning and analysis. E.J.M., B.K.L., and R.C.G. wrote the finalized models and visualized the data. E.J.M. prepared the initial draft of the manuscript. J.D.M. conceived and supervised the project, and edited the manuscript. J.D.M. and E.J.M. acquired funding for the project. All authors discussed and made contributions to the final version.

## Data and materials availability

All data and code are available in the supplementary materials.

