## Supplementary material for "Drawing the Borders of Olfactory Space": Methods and supplemental text

To build models that can predict odorous/odorless status for any molecule, we generated a large dataset with sufficient chemical diversity to learn generalizable rules relating chemical structure to odor status. We used two strategies to gather information on odor status: scraping odor publicly available classifications from the literature and websites (low cost, lower confidence) and testing compounds with human subjects (high cost, higher confidence). We classified 128 molecules as odorous or odorless through human subject testing and 1799 additional molecules through web and literature searches; in total, our dataset includes 1927 unique molecules. We used chemoinformatic softwares OpenBabel, Dragon, and EPISuite to generate chemical features for each molecule. More detailed information on the construction of this dataset follows. Code used to generate models and figures can be found at https://github.com/emayhew/OlfactorySpace.

Gathering odor classifications from literature sources and websites

Molecules with stated odorous/odorless classifications were gathered from the literature[1–6] and from databases of odorous molecules (Sigma Aldrich Flavors & Fragrances, www.sigmaaldrich.com/industries/flavors-and-fragrances.html; The Good Scents Company database, www.thegoodscentscompany.com). Additionally, we collected and classified molecules with odor information from the websites Available Chemicals Directory (psds.ac.uk/acd; keyword: odorless), Wikipedia (wikipedia.org; keyword: odorless), and PubChem (pubchem.ncbi.nlm.nih.gov; keywords: odor, odour, smell, fragrance, aroma, sense of smell, no odor, no odour, no smell, no fragrance, odorless, odourless). The source of the odorous/odorless classification for each molecule in our dataset is given in Supplementary Table 1.

### Selecting compounds for human psychophysics testing

In selecting chemical compounds to be evaluated by human subjects, we were constrained by the availability of safety data. We gathered candidate molecules from four chemical libraries in which the majority of compounds are considered safe by experts: Generally Recognized as Safe database (GRAS, www.fda.gov/food/generally-recognized-safe-gras/gras-substances-scogs-database), Prestwick chemical library (www.prestwickchemical.com/screening-libraries/prestwick-chemical-library/), The Good Scents Company database (www.thegoodscentscompany.com), and Arctander's Perfume and Flavor Chemicals database (Arctander 1969). From these databases, we gathered a total set of 6009 unique compounds and calculated structural and physicochemical descriptors (Dragon v6, Talete; www.talete.mi.it).

Machine learning (ML) models can only learn rules and patterns present in the training set. In order to build well-performing and generalizable models, it is critical to generate a training set that represents the full space that the model should describe and is drawn from the same population as the validation and test sets. To choose a set of compounds that can best represent the totality of chemical space, we performed k-means clustering on the centered and scaled Dragon descriptors with thirty clusters and aimed to pick four compounds from each cluster. By including molecules from every cluster in the training, validation, and test sets, we ensured that patterns learned by our models applied to the full range of chemical compounds and could be tested on validation and test sets with a similar composition. A total of 128 compounds (Supplementary Table 2) were selected to span the maximum range of chemical space regardless of their perceptual properties. All 128 compounds evaluated by human subjects for our study have no known risk to humans at the delivered concentrations, and have been approved by authoritative agencies such as the Food and Drug Administration (FDA) and the European Medicines Agency (EMEA).

Determining odor classification using human psychophysics

We prepared samples for human subject testing by aliquoting neat material (1 mL of liquid compounds or 1 g of solid compounds) into triple-washed amber jars (2 acidic and 1 neutral wash cycle). Compounds with an extremely strong odor or compounds that were not available in large quantities (e.g. reserpine) were presented at lower concentrations or in dilution; information on compound dosing is included in Supplementary Table 2. Between experimental sessions, we stored jars containing chemical compounds in a 3-shelf acrylic cabinet supplied by a low flow of continuous carbon-filtered air to ensure the stored odorants would not be contaminated by outside ambient air or other jars. During testing sessions, experimenters wore lab coats and odorless gloves to prevent contamination of the jars.

Ninety unique normosmic participants (64 females) aged 18-54, (median 27.9) who consented to our study procedure previously approved by University of Pennsylvania Institutional Review Board were tested. Approximately 15 participants were recruited for each set of 5 compounds (hereby referred to as “blocks”) but were permitted to participate in multiple blocks. Twenty-six blocks were run in total (130 compounds).

Each test session consisted of 28 trials: 5 target compounds were presented 5 times each alongside two blank jars in a 3-alternative forced choice test. Participants were asked to choose the jar that contained an odorant, as well as rate their confidence in choosing the correct jar, ranging from 0 (Completely Unsure) to 10 (Extremely Confident). The remaining 3 trials were presentations of a control compound, linalool (1 mL, neat), to screen participant olfactory function. All participants were blindfolded during the entire test session, and the jars were delivered to the nose of participants by the experimenter. Participants also performed an auditory task during 30-second breaks between each trial to maintain subject attention while minimizing olfactory adaptation. The order of presentations of target and blank jars were randomized and responses were recorded using E-prime 3.0 (Psychology Software Tools, Pittsburgh, PA). The University of Pennsylvania Institutional Review Board approved this research protocol, and all human research participants gave informed consent.

After each block was completed, the probability of detection of each compound within the block was calculated. The scale of probability ranges from p=0.0 (undetectable) to p=1.0 (perfect detection), with a chance detection probability of p=0.33. Based on the binomial probability distribution for n=75 trials and chance p=0.33 (the cumulative, one-tailed probability of recording 33 or more correct trials out of 75 is p=0.035), compounds correctly discriminated from blanks in ≥ 33 trials were determined to be odorous (α=0.05). Once ≥ 33 correct identifications were made for all compounds in a block, we terminated data collection for that set of compounds and moved on to the next block. The number of completed and correct trials for each compound is included in Supplementary Table 1, and the proportion of correct trials for each molecule is plotted in Extended Data Fig. 3.

<u>Correcting for odorous impurities in odorless compounds</u>

We performed headspace extraction using StableFlex 2CM solid-phase microextraction (SPME) fibers (Supelco), exposing the SPME fibers for 5 minutes to the headspace of a jar prepared as described above. We then inserted the SPME fibers (60 s desorption time) into Thermo Scientific ISQ single-stage quadrupole GC-MS and Thermo-Fisher Trace GC Ultra GC-O instruments fitted with identical Restek 30-meter Stabilwax columns (1 μm coating) so that a given molecule should elute with the same retention time (RT) on both instruments. We used Xcalibur software (Thermo Electron Corp.) to analyze the GC-MS spectra and identified all compounds present above 10% relative abundance. In the case of a few compounds for which the volatility was too low for SPME, we performed direct injection of analytes (100 ppm). In direct injection experiments, the most abundant compound is guaranteed to be the nominal compound, and RT for the target molecule was determined by FID. We recorded the RT and odor quality notes for all odors perceived during each GC-O experiment.

Compounds which had either no peak in the GC-MS spectra or no corresponding GC-O odor within 60s of the GC-MS(RT) were reclassified as odorless. In total, 23 of the 111 compounds tested (21%) were reclassified, highlighting the importance of controlling for impurities in olfactory research. Results of QC on odor classification are shown in Extended Data Fig. 3, and information on the cause of GC reclassifications is included in Supplementary Table 2.

Preparing data for model-building

We pooled data on 128 compounds tested in our human psychophysics experiment with 1796 additional molecules for which an odor classification was available from literature sources or websites. Of these 1924 molecules, 1615 were classified as odorous and 309 were classified as odorless. We used Open Babel[7] to generate an energy-minimized 3-dimensional structure file (.sdf) for each molecule from the Simplified Molecular-Input Line-Entry System (SMILES) string. Next, we used the chemoinformatic software Dragon (Talete v6; http://www.talete.mi.it) to calculate 4885 structural and physicochemical descriptors based on the 3-D structure. Additionally, we gathered data on the boiling point and vapor pressure of the molecules. We calculated an approximate boiling point for each molecule using two published methods[8,9]. Due to large observed errors between experimental values and estimates generated with these methods, we also generated estimated and experimental boiling point and vapor pressure values using the program EPI Suite (U.S. EPA). Finally, we collected experimental boiling point values from PubChem. We used experimental data in all cases where it was available and estimates only where it was not. Boiling point estimates calculated using the Banks method were used only for comparison with experimental data and to make predictions on GDB molecules where no experimental values were available. Importantly, while we used multiple sources to generate chemical descriptors, the three features used by our transport ML model (boiling point, vapor pressure, octanol/water partition coefficient) are all available through open-access sources (EPI Suite, U.S. EPA).

Prior to any data analysis or modeling, we removed 60 compounds from the data set to form a test set and validation set of 30 compounds each (Supplementary Table 1). Previous studies show that while ML approaches can learn well from noisy training sets, it is critical to have high confidence in the classification of examples used to measure model performance 10. Both the validation and test set molecules were drawn exclusively from the pool of 128 molecules tested in the lab. Because we wanted to ensure that validation and test sets were representative of the full data set, both datasets were composed of 1 molecule from each of 30 clusters and had an odorous:odorless ratio of 80:20 (the full lab-tested data set ratio at that time). The remaining 1867 compounds (68 tested in lab, 1796 gathered from the literature and websites) were pooled to form our training set (Supplementary Table 1). Once the model training parameters were finalized, the 30 validation molecules were added to the training set.

We dropped any features with more than 10% missing values and then performed k-nearest neighbor imputation for remaining missing values (k=5, R package bnstruct v1.0.8). We normalized the raw physicochemical descriptors in the training set by scaling each descriptor from 0 to 1, then applied the same normalization factors to descriptors in the validation and test sets. Consequently, the model was blind to the range of descriptors outside of the training set[11,12]. We then trimmed the dataset by dropping descriptors that were highly correlated (r > 0.99) or had negligible variation between molecules. All data processing and analysis was conducted using the open-source statistical software R (version 3.5.3), and preprocessing steps were done using the preProcess function in the R package caret (version 6.0.81)[13].

Training and evaluating models

Selecting the best algorithm for a problem is often an empirical process. We compared the performance of four ML algorithms that have been successfully applied in related research to generate odor classification models: support vector machine (SVM)[14,15], random forest (RF)[16–18], stochastic gradient boosting (GB)[19,20], and extreme gradient boosting (XGB)[21,22]. Models were optimized during training to maximize the area under the receiver operating characteristic curve (AUROC) on cross-validation splits. We chose to optimize models based on AUROC because it more heavily penalizes false positives, making it a better metric than accuracy in cases with imbalanced classes. All models were trained using the caret package for R 13 (training control parameters: method = repeatedcv, number = 5, repeats = 2). We addressed this significant class imbalance in our dataset (1619 odorous:310 odorless) by applying a Synthetic Minority Over-sampling TEchnique (SMOTE) to our training set using the R package DMwR (version 0.4.1)[23]. We tested several ratios of odorous:odorless training sets and chose the ratio (62 odorous:38 odorless) which optimized cross-validation AUROC.

The XGB algorithm produced the best-performing models as assessed through cross-validation AUROC and was chosen to generate the final ML models.

Comparing model performance across chemical classes

To evaluate the performance of our models across chemical classes, we generated 80 randomized train-test splits (80:20 ratio) from our full dataset of 1924 molecules. We trained each model and evaluated its AUROC performance on the relevant subset of the test split. We report average AUROCs across the 80 randomized train-test splits. Each chemical class subset was constructed by filtering for molecules matching the SMARTS queries below.

| **Chemical class** | **SMARTS query used** |
|---|---|
| Benzene | c1ccccc1 |
| Ester | [CX3](=O)O[#6] |
| Carboxylic Acid | [CX3](=O)[OX2H1] |
| Aldehyde or ketone | [$([#6][CX3](=O)[#6]),$([#6][CX3H1](=O))] |
| Alkyl (4+ carbon chain) | [CH2X4][CH2X4][CH2X4][CH2X4] |
| Amine | [NX3;H2,H1,H0;!$(NC=O);!$(NO)] |
| Organohalide | [#6][#9,#17,#35,#53] |
| Hydroxyl | [C;!$(C=O)][OX2H] |
| Ether | [!$(C=O);$([C&!a])][OX2&H0][!$(C=O);#6] |
| Inorganic | [#6] (Query results were inverted as SMARTS does not support negative matches) |
| Organosulfide | [SX2][#6] |

Estimating number of possible odorants

In order to generate an estimate of the total number of possible odorants, we applied an XGB-trained transport model based on estimated boiling point and log P to a representative set of molecules from the GDB chemical database[24,25]. Estimated boiling point values were used because experimental values are not available for the majority of molecules in the GDB database. The database was binned by heavy atom count (HAC), and each bin was randomly downsampled to contain no more than 10,000 molecules. The final GDB subset included 107,086 total molecules (Supplementary Table 3).

To estimate the possible number of odorous molecules, we need to estimate both the possible number of unique molecules and the proportion of those molecules that will be odorous. The GDB-17 database generated by Ruddigkeit et al[25] includes 166.4 billion molecules with 17 or fewer heavy atoms composed of only C, H, N, O, S, and halogens; this dataset provides a conservative estimate of the number of possible, relevant molecules. In a later study, Ruddigkeit et al[26] found that known fragrance molecules have up to 21 heavy atoms. The number of possible molecules is difficult to estimate, but we project that molecules with HAC > 17 will have at least as many molecules as HAC 17, of which there are 109,481,780,580 unique molecules in GDB-17.

The proportion of molecules predicted to be odorous is heavily dependent on HAC. We calculated an average odorous probability for each HAC from 1 to 17. In our conservative estimate, we simply multiplied the average odorous probability by the number of molecules of that HAC in GDB-17 and summed these values across HAC. We subsequently used transport model-predicted odorous probabilities to generate 10 binary odorous/odorless classifications for each of the 107,086 molecules in our GDB subset. The standard error bars in Fig. 3a represents the variation in the proportion of odorous molecules by HAC across the 10 simulations. We then fit a logistic regression model (glm function, R) to the binary odor classifications as a function of HAC. We used this model to extend our estimates of the proportion of molecules that are odorous to HAC 21. Our less conservative estimate of possible odorous molecules assumes HAC 18-21 have at least 109 billion molecules each and is calculated by multiplying GDB molecular count estimates by the logistic regression model proportion odorous estimates for each HAC up to HAC 21.

Because our model is based on principles of physical transport which apply to all molecules, we anticipate that our model will produce accurate predictions for all GDB-17 molecules. However, to provide a more cautious estimate, we also segmented our down-sampled set of GDB-17 molecules based on the distance to their nearest neighbor in our training data (Tanimoto similarity). In extended Data Fig. 2, we show the cumulative number of predicted novel odorants as a function of ML Transport model-generated odorous probabilities for several Tanimoto similarity cut-offs. Considering only GDB-17 molecules with structurally similar molecules in our training data (Tanimoto similarity > 0.4), we calculate a lower bound estimate of 10 million predicted odorants in GDB-17 with structurally similar known odorants.

Our estimate is sensitive to the composition of our training set: increasing the ratio of odorous molecules in the dataset increases the proportion of predicted odors. However, the poor model performance with extreme training set imbalance suggests that balanced training sets like ours gives better estimates of size of odor space.

**Visualizing odors in chemical space**

A two-dimensional semi-supervised embedding of odor space was created by applying the UMAP algorithm[27] to a random sample (n=107,086 molecules) of the GDB-17 database along with an additional n=8,366 molecules drawn from the experimental olfaction literature. All molecules containing atoms other than H, C, Cl, F, I, N, O, S, P (n=420 molecules) were excluded. N=4,864 physicochemical features were computed using Dragon 6.0 software. Missing feature values (0.06%) were median imputed and then all features were scaled between 0 and 1. The UMAP algorithm was also shown the model-predicted odorous probabilities of a random 25% of the odorants to assist in clustering molecules of similar odor probability. An interactive version of is available at https://odormap.pyrfume.org.

**Data and materials availability:** All code and data needed to train the models and reproduce the analyses shown in Figures 1-2 will be publicly available at the URL https://github.com/emayhew/OlfactorySpace following publication of the study. There are no restrictions on the availability of this data. Due to licensing restrictions, we are not providing Dragon descriptors for the full GDB (Generated DataBase) or common flavor and fragrance compounds.

**Code availability:** All code needed reproduce the analyses described in this paper are publicly available at the URL https://github.com/emayhew/OlfactorySpace. There are no restrictions on the availability of this code.

**Acknowledgements**

The authors acknowledge support from the George Preti Research Support Core for Analytical Chemistry and would like to thank George Preti, Katherine A. Prokop-Prigge, Bruce A. Kimball, Kai Zhao, and Gary K. Beauchamp for comments and discussions. This research was supported in part by grants from the NIH (R01DC013339, U19NS112953, R01DC018455, and

R01DC017757) and the Ajinomoto Co. Innovation Alliance Program. E.J.M. was supported by T32DC000014 and F32DC019030.

**Author contributions**

C.J.A., J.M.M., L.L.S., and C.W.Y. collected human behavioral data. L.L.S., K.A.L., and E.J.M carried out quality control using GC-O. E.J.M., C.W.Y., K.A.L., R.C.G., and J.D.M. curated external data. B.K.L. validated the models and analyzed the domain applicability. C.W.Y and C.J.A conducted initial data cleaning and analysis. E.J.M, B.K.L., and R.C.G. wrote the finalized models and visualized the data. E.J.M. prepared the initial draft of the manuscript. J.D.M. conceived and supervised the project, and edited the manuscript. J.D.M. and E.J.M. acquired funding for the project. All authors discussed and made contributions to the final version.

**Competing interest declaration**

J.D.M. received research funding from Ajinomoto Co., Inc.

**Extended data figures**

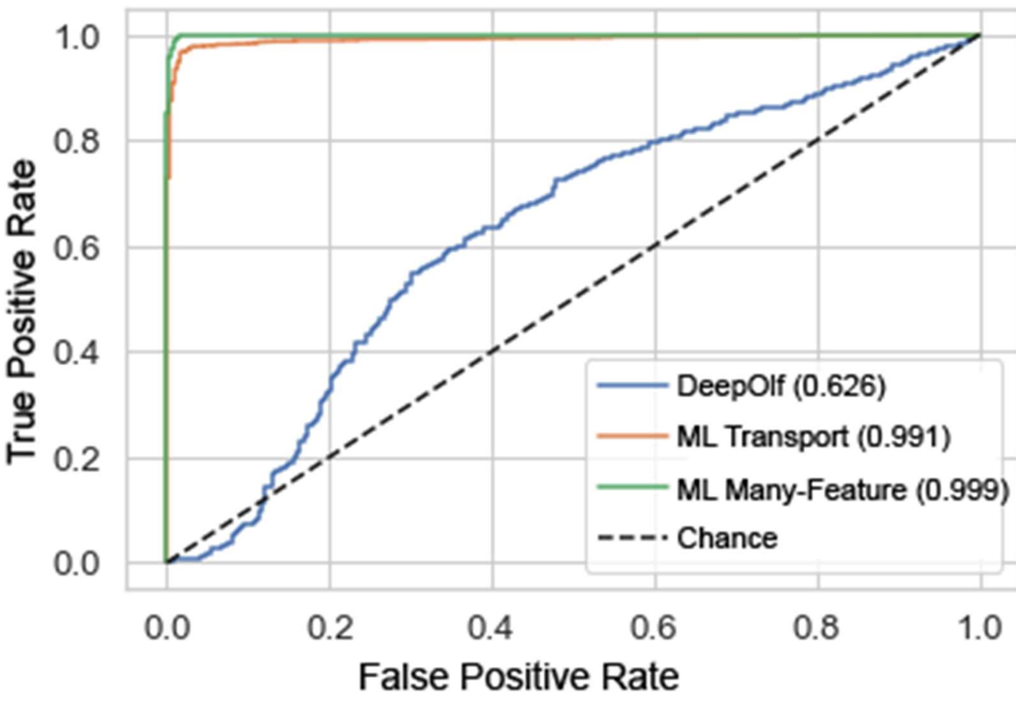


**Extended Data Figure 1. DeepOlf model underperforms on cleaned dataset.** We measured the performance of the DeepOlf model published by Sharma et al (2020)[6] on our full cleaned dataset (n=1924); this plot of true positive rate versus false positive rate shows the receiver operating characteristic (ROC) curves for the DeepOlf, ML Transport, and ML Many-Feature models. Chance classification accuracy is represented as a dotted black line. The area under the ROC curve for each model is reported in parentheses next to the model name.

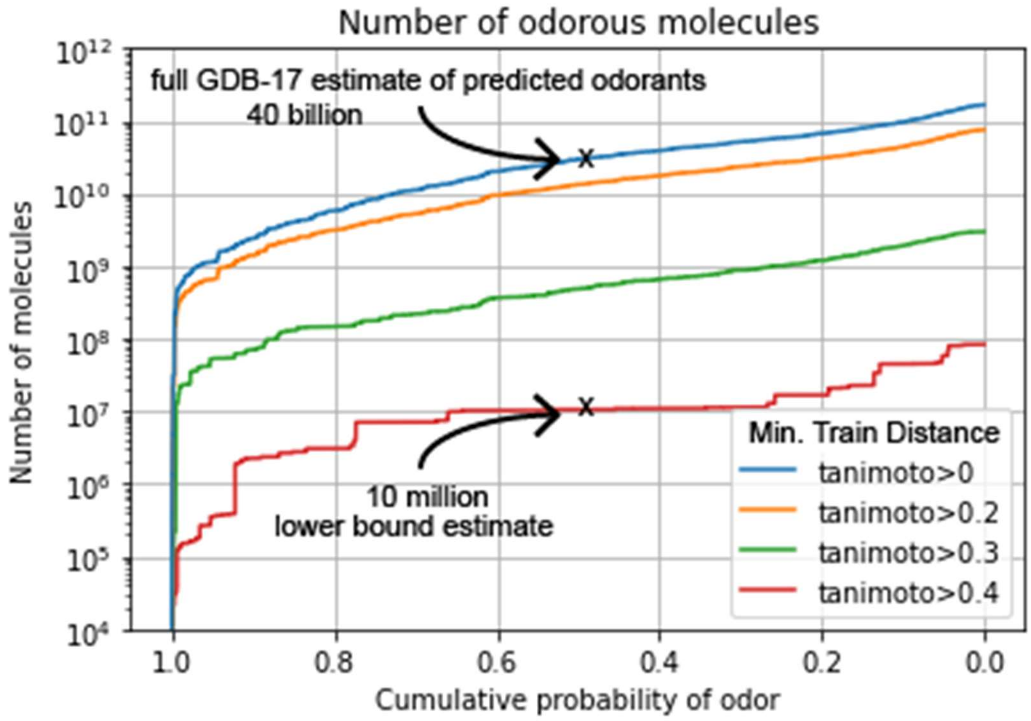


**Extended Data Figure 2. Possible odorant estimates as a function of nearest training set example.** Our transport model predicts 40 billion molecules in GDB-17 will be odorous (probability of having an odor > 0.5). By segmenting GDB-17 molecules by the minimum Tanimoto distance to a training set molecule ("Min. Train Distance"), we show a conservative lower bound estimate of 10 million predicted odorous molecules that have a structurally similar neighbor (Tanimoto similarity > 0.4) in our training data.

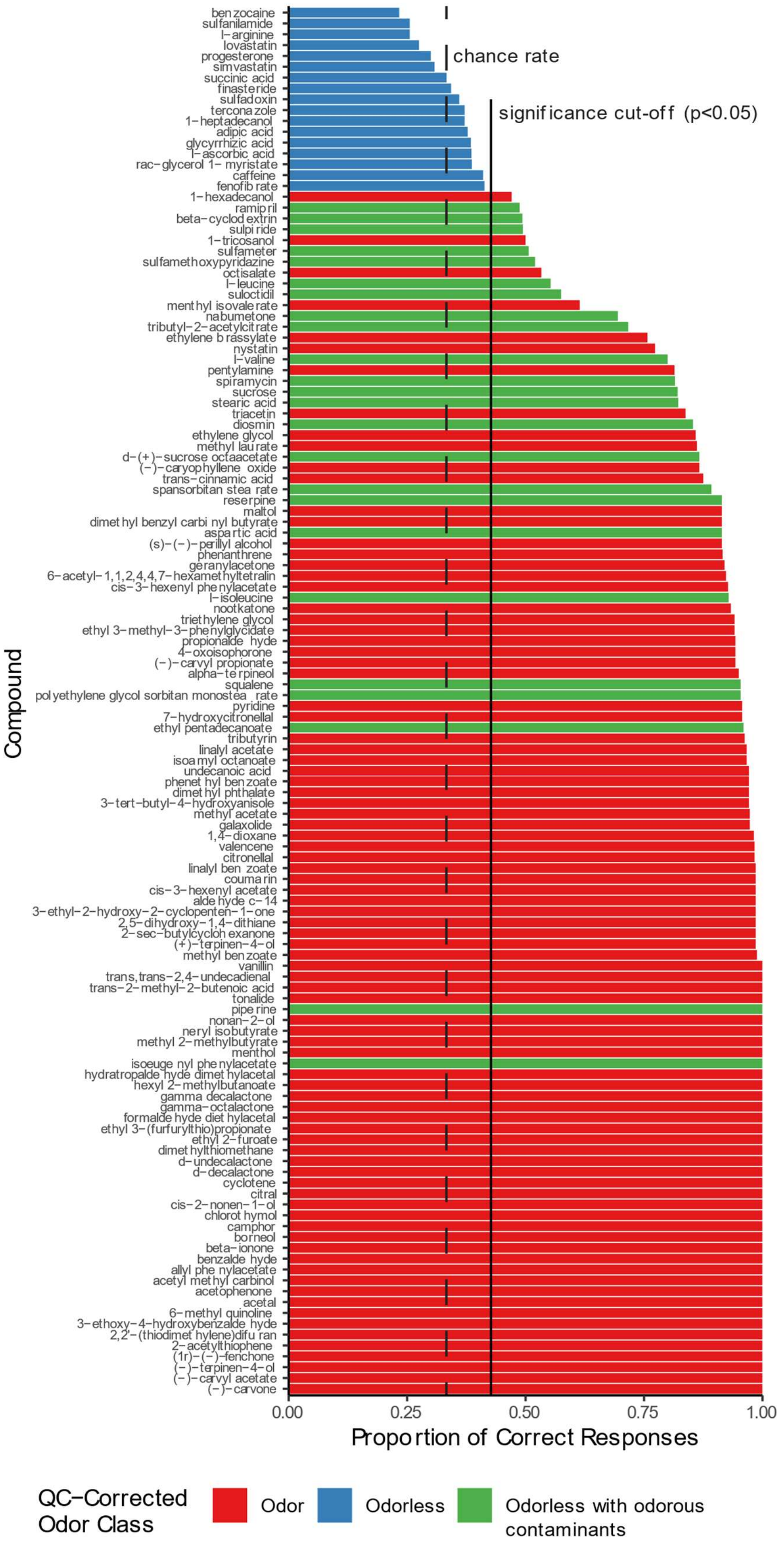

benzocaine
sulfanilamide
l−arginine
lovastatin
progesterone
simvastatin
succinic acid
finasteride
sulfadoxin
terconazole
1−heptadecanol
adipic acid
glycyrrhizic acid
l−ascorbic acid
rac−glycerol 1− myristate
caffeine
fenofibrate
1−hexadecanol
ramipril
beta−cyclodextrin
sulpiride
1−tricosanol
sulfameter
sulfamethoxypyridazine
octisalate
l−leucine
suloctidil
menthyl isovalerate
nabumetone
tributyl−2−acetylcitrate
ethylene brassylate
nystatin
l−valine
pentylamine
spiramycin
sucrose
stearic acid
triacetin
diosmin
ethylene glycol
methyl laurate
d−(+)−sucrose octaacetate
(−)−caryophyllene oxide
trans−cinnamic acid
spansorbitan stearate
reserpine
maltol
dimethyl benzyl carbinyl butyrate
aspartic acid
(s)−(−)−perillyl alcohol
phenanthrene
geranylacetone
6−acetyl−1,1,2,4,4,7−hexamethyltetralin
cis−3−hexenyl phenylacetate
l−isoleucine
nootkatone
triethylene glycol
ethyl 3−methyl−3−phenylglycidate
propionaldehyde
4−oxoisophorone
(−)−carvyl propionate
alpha−terpineol
squalene
polyethylene glycol sorbitan monostearate
pyridine
7−hydroxycitronellal
ethyl pentadecanoate
tributyrin
linalyl acetate
isoamyl octanoate
undecanoic acid
phenethyl benzoate
dimethyl phthalate
3−tert−butyl−4−hydroxyanisole
methyl acetate
galaxolide
1,4−dioxane
valencene
citronellal
linalyl benzoate
coumarin
cis−3−hexenyl acetate
aldehyde c−14
3−ethyl−2−hydroxy−2−cyclopenten−1−one
2,5−dihydroxy−1,4−dithiane
2−sec−butylcyclohexanone
(+)−terpinen−4−ol
methyl benzoate
vanillin
trans,trans−2,4−undecadienal
trans−2−methyl−2−butenoic acid
tonalide
piperine
nonan−2−ol
neryl isobutyrate
methyl 2−methylbutyrate
menthol
isoeugenyl phenylacetate
hydratropaldehyde dimethylacetal
hexyl 2−methylbutanoate
gamma decalactone
gamma−octalactone
formaldehyde diethylacetal
ethyl 3−(furfurylthio)propionate
ethyl 2−furoate
dimethylthiomethane
d−undecalactone
d−decalactone
cyclotene
citral
cis−2−nonen−1−ol
chlorothymol
camphor
borneol
beta−ionone
benzaldehyde
allyl phenylacetate
acetyl methyl carbinol
acetophenone
acetal
6−methyl quinoline
3−ethoxy−4−hydroxybenzaldehyde
2,2'−(thiodimethylene)difuran
2−acetylthiophene
(1r)−(−)−fenchone
(−)−terpinen−4−ol
(−)−carvyl acetate
(−)−carvone
chance rate
significance cut-off (p<0.05)
Compound
0.00
0.25
0.50
0.75
1.00
Proportion of Correct Responses
QC−Corrected
Odor Class
Odor
Odorless
Odorless with odorous
contaminants

**Extended Data Figure 3. Classification of molecules as odorous or odorless through human psychophysics experiments and analytical quality control.** Blindfolded subjects were presented with 1 compound-containing jar and 2 blank jars in each trial of a 3-alternative forced choice task; the proportion of correct responses is plotted for the 128 lab-tested molecules The chance rate of selecting the correct jar is ⅓, and the minimum statistically significant (a = 0.05) correct selection rate is 0.43 (> 32 correct selections). Molecules correctly differentiated from the blanks in more than 32 trials were initially classified as odorous, plotted in red or green, while molecules correctly selected below this rate were classified as odorless, plotted in blue. A paired gas chromatography-mass spectrometry/gas chromatography-olfactometry (GC-MS/GC-O) quality control (QC) procedure was applied to identify cases in which odorous contaminants, and not the nominal compound, were responsible for the human-detectable odor; these molecules were reclassified as odorless (plotted in green).

**Methods References**


1. Boelens, H. Structure-activity relationships in chemoreception by human olfaction. *Trends Pharmacol. Sci.* **4**, 421–426 (1983).

2. Abraham, M. H., Sánchez-moreno, R., Cometto-muñiz, J. E. & Cain, W. S. An algorithm for 353 odor detection thresholds in humans. *Chem. Senses* **37**, 207–218 (2012).

3. Hau, K. M. & Connell, D. W. Quantitative structure-activity relationships (QSARs) for odor thresholds of volatile organic compounds (VOCs). *Indoor Air* **8**, 23–33 (1998).

4. Arctander, S. *Perfume and Flavor Chemicals*. (Allured Publishing Corporation, 1969).

5. Laffort, P. & Gortan, C. Olfactory properties of some gases in hyperbaric atmosphere. *Chem. Senses* **12**, 139–142 (1987).

6. van Gemert, L. J. *Odour Thresholds*. (Oliemans Punter & Partners BV, 2011).
